# Immune potentiator for increased safety and improved protection of vaccines by NF-kB modulation

**DOI:** 10.1101/489732

**Authors:** Brittany A. Moser, Rachel C. Steinhardt, Yoseline Escalante-Buendia, David A. Boltz, Kaylynn M. Barker, Stan Yoo, Bethany G. McGonnigal, Aaron P. Esser-Kahn

**Affiliations:** Institute for Molecular Engineering, University of Chicago, 5640 South Ellis Avenue, Chicago, Illinois 60637, USA; Division of Microbiology and Molecular Biology, IIT Research Institute, Illinois Institute of Technology, 10W. 35th Street, Chicago, IL 60616, USA; Department of Chemistry, Chemical Engineering & Materials Science, Biomedical Engineering, University of California, Irvine, California 92697, USA

## Abstract

Many modern vaccines include adjuvants that activate the immune system and provide an enhanced humoral or cellular response. Current approved adjuvants are unable to provide desired responses against some pathogens (e.g. HIV or dengue). Many new adjuvants have been developed and demonstrate promising results, but side effects from the inflammatory response induced by these adjuvants have resulted in limited FDA approvals. No adjuvants yet possess the capability to independently modulate inflammation and protection. Here we demonstrate a method to limit inflammation and side effects associated with vaccination while retaining the protective responses using a variety of promising adjuvants. To accomplish this, we combined a selective NF-kB inhibitor with the immune adjuvant. The resulting vaccines reduce systemic inflammation and boost antibody responses. In an influenza challenge model, we demonstrate that this approach enhances protection. This method is generalizable across a broad range of adjuvants and antigens. We anticipate these studies will lead to a novel approach to vaccine formulation design that may prove general across a wide range of adjuvants, enabling their greater use in the public realm.

## Main Text

Vaccines are considered one of the most effective global health interventions against infectious diseases. Despite their success, current and future vaccines face contradictory challenges of increasingly stringent safety margins and more effective and diverse protective responses. A major challenge in developing new vaccine approaches is striking a balance between effective immune activation, leading to protective responses, and limiting the excess inflammation and side effects. To boost the immune response, toll-like receptor (TLR) agonists have been explored as vaccine adjuvants because they activate the innate immune system, promoting the expression of inflammatory cytokines and cell surface receptors important for T cell interactions (*1–6*). Effective TLR agonists stimulate the desired cellular or humoral adaptive responses; however, the inflammation induced by many of these compounds has made it challenging to transition them into new clinical vaccines (*7*). For example, CpG DNA, a TLR 9 agonist, has wide-ranging promise as a vaccine adjuvant and provides protection for diseases currently without a vaccine, such as HIV (*8*). CpG DNA also enables vaccines to be produced with less antigen (*9*), induces protective responses faster (*10*), and produces effective anti-tumor activity (*11,12*). However, the excessive inflammatory response induced by this adjuvant has resulted in many clinical trial failures and is cited as limiting its therapeutic promise (*13,14*). CpGs are only a fraction of the hundreds of TLR agonists (*15*). However due to the unsafe side effects, only a handful of TLR agonists are approved for limited use in humans (*16*). Studies indicate that side effects are mediated through systemic distribution of TNF-a and IL-6 (*17,18*). Here we demonstrate a method to decouple part of the inflammatory response from the antigen presenting actions of several adjuvants using an immune potentiator. Using a broad range of TLR agonists, we demonstrate both in vitro and in vivo that using an immune potentiator decreases proinflammatory cytokines while maintaining adaptive immune function. In vivo, we find that co-administering the immune potentiator with the 2017-2018 flu vaccine (Fluzone) decreases side effects associated with vaccination and increases protection. Co-administration of the immune potentiator with CpG-ODN1826 (CpG) and dengue capsid protein leads to elimination of systemic proinflammatory cytokines post-vaccination and yields increased, neutralizing antibodies. Additionally, administering the immune potentiator with CpG and gp120, a HIV viral coat protein, increased serum IgG and vaginal IgA antibodies and shifted IgG antibody epitope recognition. Lastly, we observed immune potentiation for several TLR agonists – implying a general approach. Immune-potentiation may find use in reducing the systemic side effects associated with inflammation for many adjuvanted vaccines (*19*) – creating the potential for many PRR agonists to be used safely, increasing the diversity of adaptive immune profiles and widening the scope of disease prevention and treatment.

## Selection of Immune Potentiator

In seeking a method of immune potentiation, we explored the extensive research on the TLR activation pathway. This powerful mechanistic framework let us hypothesize about how TLR activation directs inflammatory cytokines and antigen presentation. As TLR pathways converge with NF-kB activation, and inflammatory and adaptive responses diverge upon which NF-kB subunit is activated, we hypothesized that we could decouple these processes via selective inhibition – leading to reduced side effects but maintaining the adaptive response. Upon TLR activation, the transcription factor NF-kB primes the transcription of pro-inflammatory cytokines such as IL-6 and TNF-a, and cell surface receptors such as MHC-II, CD40, CD80 and CD86 (*20–22*). The NF-kB family is a family of transcription factors, consisting of two subunits: a DNA binding domain and a transcriptional activator (*23,24*). Each NF-kB dimer controls expression of a different set of genes for distinct cellular processes – broadly, some dimers control inflammatory expression while others control antigen presentation (*23–25*). Selectively modulating a pathway, we conjectured, might lead to increased antigen presentation, while decreasing inflammation. NF-kB inhibitors have been widely explored for reducing cytokine expression in cancer (*26–29*), autoimmune disorders (*30,31*), and sepsis (*32–34*), yet they have not been explored as vaccine potentiators. This lack of experimentation may be because it is broadly understood that NF-κB activation is necessary in mounting an adequate adaptive immune response (*29,35*). However, only certain subunits direct antigen presentation (*36*). As a proof-of-concept immune potentiator we chose SN50, a cell permeable peptide that consists of the nuclear localization sequence (NLS) of the NF-kB subunit, p50 which blocks the import of p50 containing dimers into the nucleus (*37*).

First, we sought to determine if SN50 enables inhibition of NF-kB of innate immune cells. We validated that SN50 reduced total NF-kB activity in human (THP-1 monocytes) and mouse (RAW macrophages) cells in a dose dependent manner. (**Fig.S1a-d**).

## Examination of CpG-induced inflammation and resulting immune response

We sought to verify that SN50 could enable antigen presenting cells to upregulate cell surface receptors, while limiting pro-inflammatory cytokine production. We incubated murine bone marrow-derived dendritic cells (BMDCs) with SN50 and CpG or CpG alone for 6h and analyzed how the potentiator altered cytokine production and cell surface receptor expression (**Fig. 1a**). Intracellular cytokine staining revealed that cells treated with SN50 demonstrated a 21% decrease in cells expressing TNF-a and a 13% decrease in cells expressing IL-6. Meanwhile, CD86 was upregulated by 22% and CD40 was only down regulated by 2.5%. Because the p65-p50 dimer is the most abundant dimer found in resting cells and involved in inflammatory cytokine production, we conjecture that by inhibiting this dimer, we enable the transcription and translation of cell surface receptors while limiting inflammatory cytokines. This is consistent with previous knockout experiments (*36*). The result is lower inflammatory responses while priming effective adaptive immune communication.

**Figure 1.**
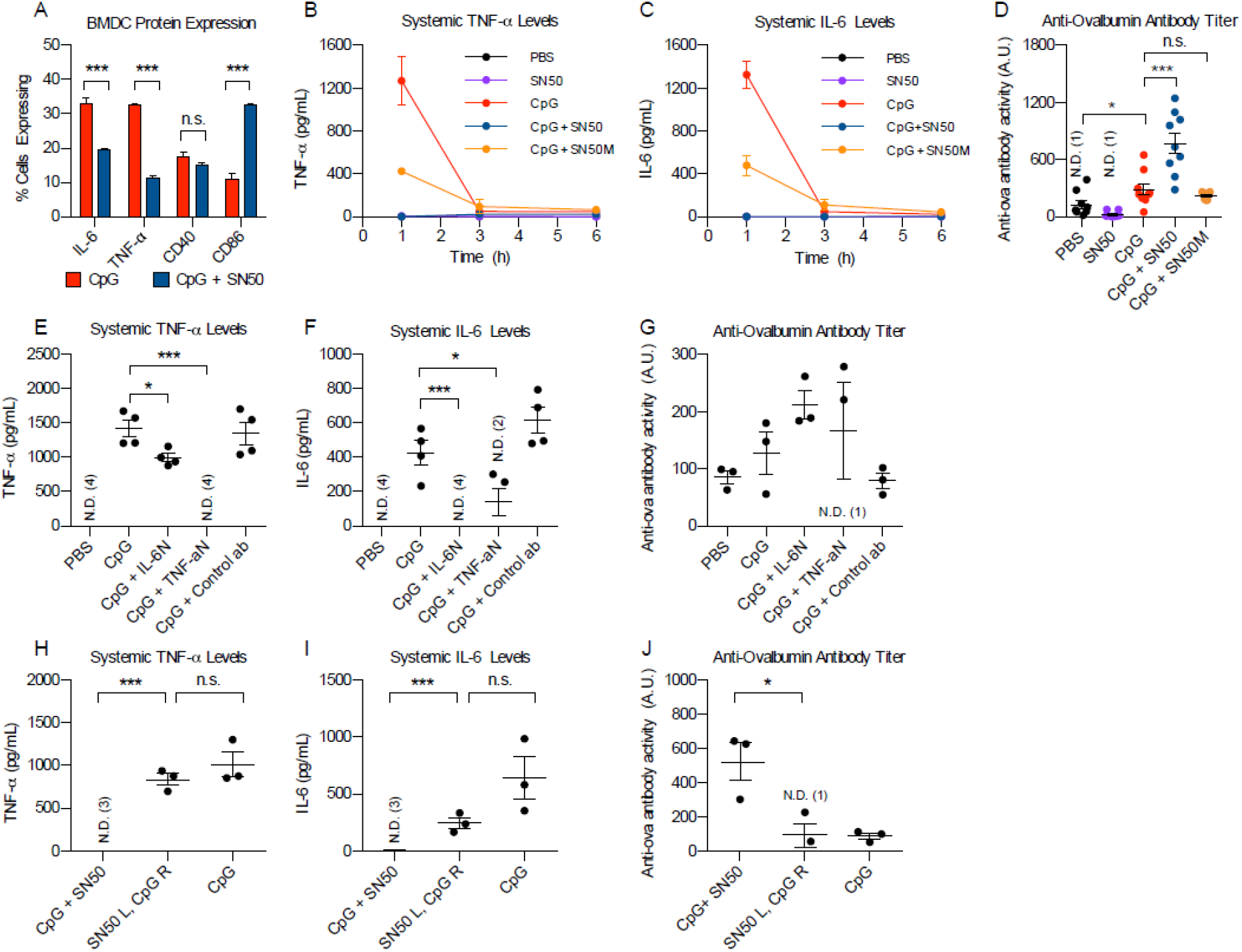
In vivo vaccination with model antigen ovalbumin and immune adjuvant SN50. (A) Intracellular cytokine staining of BMDCs treated with CpG (red bars) or CpG + SN50 (blue bars). (B) Systemic cytokine levels of TNF-a measured at 1h, 3h, 6h post-injection with: PBS (black line), SN50 alone (purple line), CpG (red line), CpG + SN50 (blue line), CpG + SN50M (yellow line), n = 4 for each time point. (C) Systemic cytokine levels of IL-6. (D) Anti-ovalbumin antibody titer, day 28, n = 8. (E) Systemic TNF-a levels 1h post-vaccination with CpG, CpG + IL-6N, CpG + TNF-aN or CpG + Control ab, n = 4. (F) Systemic IL-6 levels 1h post-vaccination with CpG, CpG + IL-6N, CpG + TNF-aN or CpG + Control ab. (G) Anti-ovalbumin antibody titer, day 28. (H) Systemic TNF-a levels in mice vaccinated with mixed CpG and SN50 (CpG + SN50), SN50 in left limb and CpG + OVA in right limb (SN50L, CpG R, or CpG alone, n =3. (I) Systemic IL-6 levels. (J) Anti-OVA antibody titer, day 28.

After observing that SN50 can limit inflammation without decreasing cell surface receptor expression in vitro, we next wanted to examine the effect in vivo. To determine if inhibition of NF-kB could decrease the systemic levels of pro-inflammatory cytokines associated with CpG vaccination, we vaccinated mice intramuscularly (i.m.) with 100 μg ovalbumin (OVA) and: PBS, SN50 (500 μg), CpG (50 μg), SN50 + CpG, or SN50M (500 μg) + CpG. SN50M is a physical control for SN50 as it is a much weaker inhibitor. We chose to measure systemic levels of proinflammatory cytokines TNF-a and IL-6 because high levels are unsafe and lead to side effects (*17,18,38*). We measured these pro-inflammatory cytokines at 1h, 3h, 6h, 24h and 48h post-injection in all groups to determine the timepoint where cytokines peak in response to CpG vaccination (**Fig. 1b, 1c, S2a, S2b**). Mice vaccinated with OVA and PBS or SN50 alone elicited no systemic cytokine response. CpG demonstrated the highest response of both TNF-a (1325 pg/mL) and IL-6 (1269 pg/mL) at the 1h timepoint. The CpG + SN50 group showed complete elimination of cytokines for both cytokines. The CpG + SN50M group showed a decrease in cytokine levels, although not as large as observed with CpG + SN50. We confirmed that this decrease in inflammatory cytokines is due to the high local inhibition of injected SN50M and not physical aggregation (**Fig. S3**). To determine how SN50 would affect the humoral response, we analyzed serum antibody levels on day 28 (**Fig. 1d**). The CpG group demonstrated a 2.4-fold increase in anti-OVA antibodies compared to PBS alone. Mice vaccinated with CpG + SN50 demonstrated a 5.9-fold increase over the PBS group and 2.7-fold increase over the CpG group. These data confirmed our hypothesis that high levels of systemic TNF-a and IL-6 can be decoupled from the humoral, adaptive immune response. We were surprised to find that addition of SN50 boosted the downstream adaptive response, leading to immune potentiation. Due to this increase in adaptive response and improved safety profile after vaccination we consider SN50 to be an immune potentiator.

## Determining mechanism of action

To more directly examine how early systemic expression of TNF-a and IL-6 impact the immediate inflammatory response and downstream adaptive response, we vaccinated mice with CpG and either TNF-a neutralizing antibody (TNF-aN) or IL-6 neutralizing antibody (IL-6N) and measured the systemic cytokines (**Fig. 1e, 1f**). The CpG + IL-6N group demonstrated a 1.4 -fold decrease in TNF-a expression and a complete reduction of systemic IL-6 expression. The CpG + TNF-aN group demonstrated complete elimination of systemic TNF-a and a 3-fold reduction of IL-6 expression. This result was confirmed by a control isotype antibody to rule out any nonspecific interactions. Although both IL-6N and TNF-aN groups demonstrated higher average antibody titer, these differences were not statistically significant (**Fig. 1g**). This indicates that reducing inflammation from CpG with the initial vaccination is not detrimental to antibody titer.

Upon observing this in vivo modulation, we sought to determine whether SN50 acts locally or systemically. To examine this mechanism, we injected SN50 i.m. in the left hind limb and immediately injected CpG + OVA (SN50 L + CpG R) in the right hind limb. There was no significant difference in systemic cytokine levels tested between CpG and the SN50 L + CpG R group (**Figure 1h, 1i**), whereas SN50 + CpG injected simultaneously demonstrated reduction of TNF-a and IL-6. On day 28, we analyzed antibody titer, reveling a 5.5 fold difference between the SN50 + CpG and the SN50 L +CpG R group (**Fig. 1j**). This demonstrates the importance of coadministration of the components and therefore indicates that SN50 is acting locally to both increase safety and protection.

From these experiments, we conclude that SN50 acts locally at the injection site to inhibit immediate cytokine production, containing inflammation before it is distributed systemically. Based on our in vitro data, we believe CpG + SN50 enables TNF-a and IL-6 production locally at reduced levels. Our in vitro data suggests that immune cells exposed to CpG + SN50 express higher levels of cell surface receptors important for antigen presentation and effective T cell activation. While our experiments confirm that SN50 reduces systemic inflammation and increases antibody titer in vivo, more in depth exploration needs to be completed to fully understand our hypothesized mechanism.

## Immune potentiation in in vivo influenza challenge model

We next wanted to focus on how SN50 might transition to a vaccine with challenge. We selected influenza vaccine as a proof-of-concept vaccination both due to its universality and the relative ease of running animal challenges with multiple parameters. We sought to determine if SN50 would reduce side effects associated with strong adjuvanticity and to see what effect this alteration on systemic cytokines would have on protection. We vaccinated mice i.m. with Fluzone^®^ quadrivalent vaccine (Fz) for the 2017/2018 influenza season, with or without CpG (50 μg) as an immune adjuvant and 500 μg SN50 (SN50 H) or 50 μg SN50 (SN50 L)) as an immune potentiator. The Fz + SN50 group demonstrated lower levels of TNF-a than Fz alone (**Fig. 2b, 2c**). Across all groups, the addition of SN50 reduced levels of TNF-a and IL-6 to levels consistent with the placebo group. To examine whether SN50 can mitigate side effects from vaccination, we analyzed the percent change in body weight 24, 48 and 72h post-vaccination (**Fig. 2d, S4**). Weight loss is the easiest and most objective measure of side effects in mice. Mice vaccinated with Fz and Fz + SN50 lost an average of 0.85% and 0.75%, respectively by the 24h timepoint. The Fz + CpG group lost an average of 5.9%. Adding SN50 H decreased the amount of weight loss to 2.4% and SN50 L to 5.1%. At 72h the Fz group were -1.1% of the starting weight whereas mice vaccinated with Fz + SN50 gained +1.5%. The Fz + CpG group lost -1.6% of the starting weight and adding SN50 H lead to a reduction in weight loss (0% change) and adding SN50 L lead to -1.3% change. Overall, mice with SN50 lost less weight than mice without SN50, demonstrating that SN50 lowered side effects associated with vaccination.

**Figure 2.**
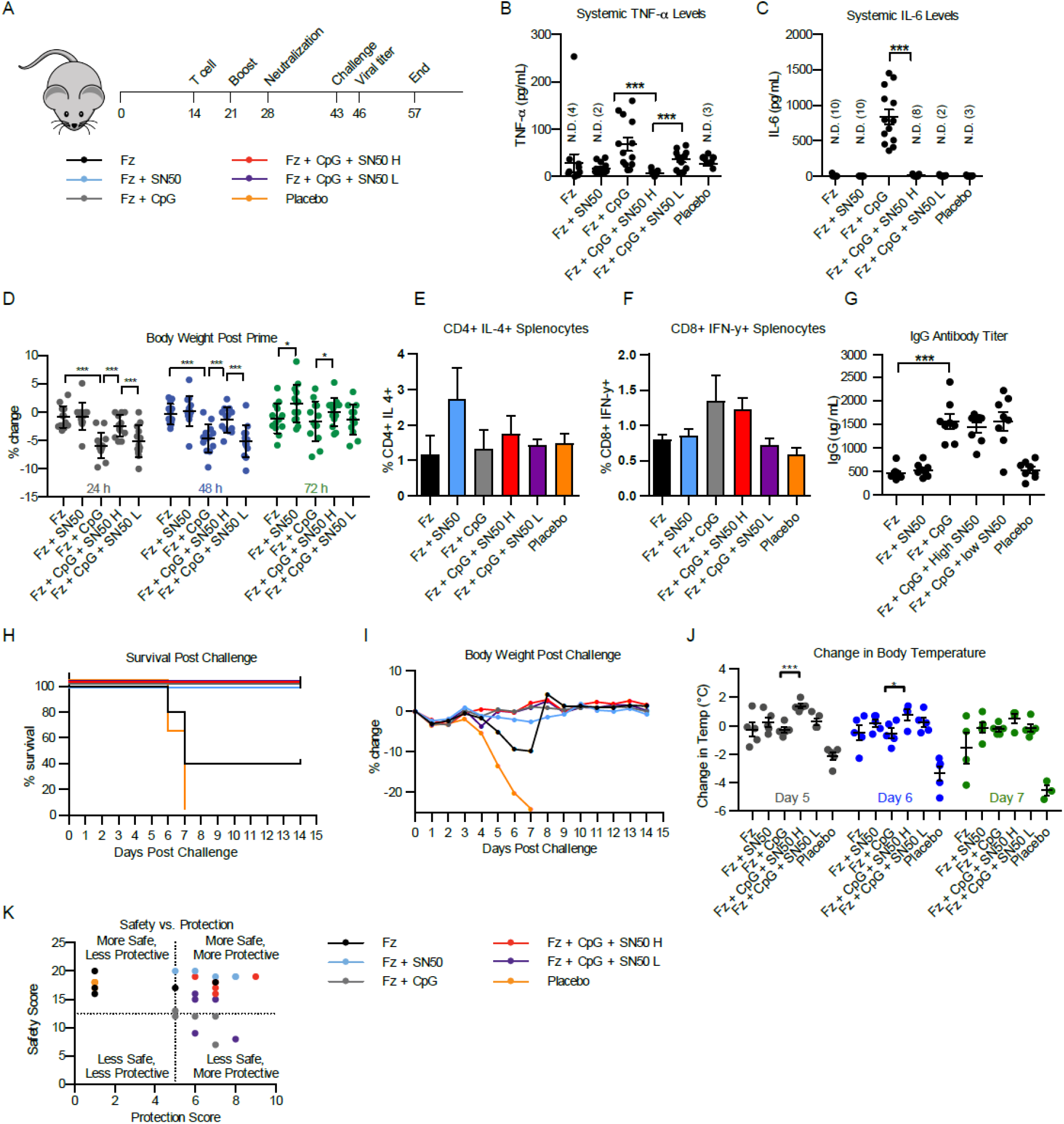
Influenza Challenge Model. (A) Schematic of influenza challenge study. (B) Systemic TNF-a levels 1h post-vaccination with Fz, Fz + SN50, Fz + CpG, Fz + CpG + SN50 H, Fz + CpG + SN50 L, Placebo. n = 13 (C) Systemic IL-6 levels 1h post-vaccination. n =13 (D) Percent change in body weight 24h (grey), 48h (blue) and 72h (green) post-vaccination, n =13. (E) Day 28 IgG antibody titer, n =8. (F) Survival 1-14 days post challenge, n = 5. Groups: Fz (black), Fz + SN50 (blue), Fz + CpG (grey), Fz + CpG + SN50 H (red), Fz + CpG + SN50 L (purple), Placebo (orange). n = 5 (G) Percent change in body weight 1-14 days, n = 5. (H) Body temperature 1-14 days post challenge, n =5. (I) Safety vs Protection score.

We next wanted to see if the addition of SN50 would change the T cell responses or antibody titer. On day 14, we analyzed splenocytes for antigen specific CD4+ and CD8+ T cells. We observed no statistically significant differences between samples with and without SN50 (**Fig. 2e, 2f**). On day 28, we analyzed the serum for antibody levels in the blood (**Fig. 2g and S5-7**). There was no significant difference in IgG titer between Fz and Fz + SN50. There was a significant difference between Fz samples and Fz + CpG of 2.9 fold. There was no significant difference between groups vaccinated with CpG, implying that the addition of SN50 reduces inflammation and side effects from vaccination, while maintaining the antibody titer.

We next sought to determine if SN50 would increase the protection of Fluzone. Mice were lethally challenged intranasally with 10^5^ PFU A/Michigan/45/2015. On day 3 post-challenge we analyzed the lungs of three mice for viral titer (**Fig. S8**). Survival was analyzed for 14 days post-challenge (**Fig. 2h**). By day 7, all placebo mice and 60% of the Fz mice had reached the humane endpoint and were euthanized. All other mice survived. The Fz + SN50 group was significantly more protected than the Fz alone group. The addition of SN50 to Fz + CpG confers equal protection, while improving side effects from the initial vaccination. Surprisingly, simply adding SN50 to Fz conferred enhanced protection equal to Fz + CpG group.

Mice were analyzed for change in body weight and body temperature for 14 days post-challenge (**Fig. 2i, 2j, Fig. S9**). The peak average weight loss between Fz (−9.9%) and Fz + SN50 (−2.67%) was statistically significant. Greater weight loss is associated with a more intense infection, these data demonstrate that adding SN50 to Fz improves the response to infection. Addition of SN50 to Fz + CpG demonstrates no significant change in weight loss indicating that the SN50 can reduce systemic cytokines and side effects from vaccination with no detrimental effects to the protective response.

As an additional parameter of disease pathology, we examined body temperature post-challenge. Unlike in humans, mice demonstrate a reduction in body temperature upon infection (*39*). The placebo has the largest peak drop in temperature (−4.57 °C), followed by the Fz group (−1.58 °C) (**Fig. 2j**). Adding SN50 to Fz or Fz + CpG mitigated the decrease in temperature across all groups.

Safety and protection of new vaccine adjuvants are typically considered two interdependent variables with an inverse relationship, where adequate protection is acquired by limiting safety or vice versa. As this potentiator makes the vaccine both safer and more protective, we sought a single way to analyze how SN50 was changing the safety and protection profile. As these variables are considered inversely related, there are few precedents for correlation. However, a common scoring system used widely across fields is a quartile-based scoring system (*40–43*). Following precedent for scoring systems, we developed a safety vs protection plot (**Fig. 2k**). This plot is meant only to serve as a visual representation of all data collected within this study. All groups that included SN50 in the vaccination increased both the safety and the protection of the vaccine. When all data is taken together, we conclude that SN50 acts as an immune potentiator by both increasing the safety profile and improving the protective outcome of the vaccination.

Next, we wanted to examine if this type of immune potentiator could improve safety and maintain the adaptive response across a broader range of diseases and antigens. We chose to vaccinate against dengue and HIV because they represent additional, important diseases with active vaccine research. In each case, challenges with current methods have been identified and we wanted to see if SN50 could help address those challenges, as well as maintain the current function of vaccination strategies. For dengue, the main challenge is producing antibodies that neutralize the virus, inhibiting cellular uptake. For HIV, a key challenge is in generating IgA antibodies at the mucosal interface as well as eliciting broadly neutralizing antibodies targeted to select epitopes. To explore how adding an immune potentiator affects each of these responses, we analyzed each antigen set in greater detail.

## Examination of immune potentiator on Dengue neutralization

To explore dengue further, we vaccinated mice with the capsid protein of dengue serotype 2 (DENV-2C) and: CpG (50 μg), CpG + 500 μg SN50. SN50 completely eliminated expression of systemic cytokines (**Fig. 3a, 3b**). On day 28 we analyzed the difference in antibody titer (**Fig. 3c**). Antibody titer in CpG + SN50 mice were almost two-fold higher than the CpG group.

**Figure 3.**
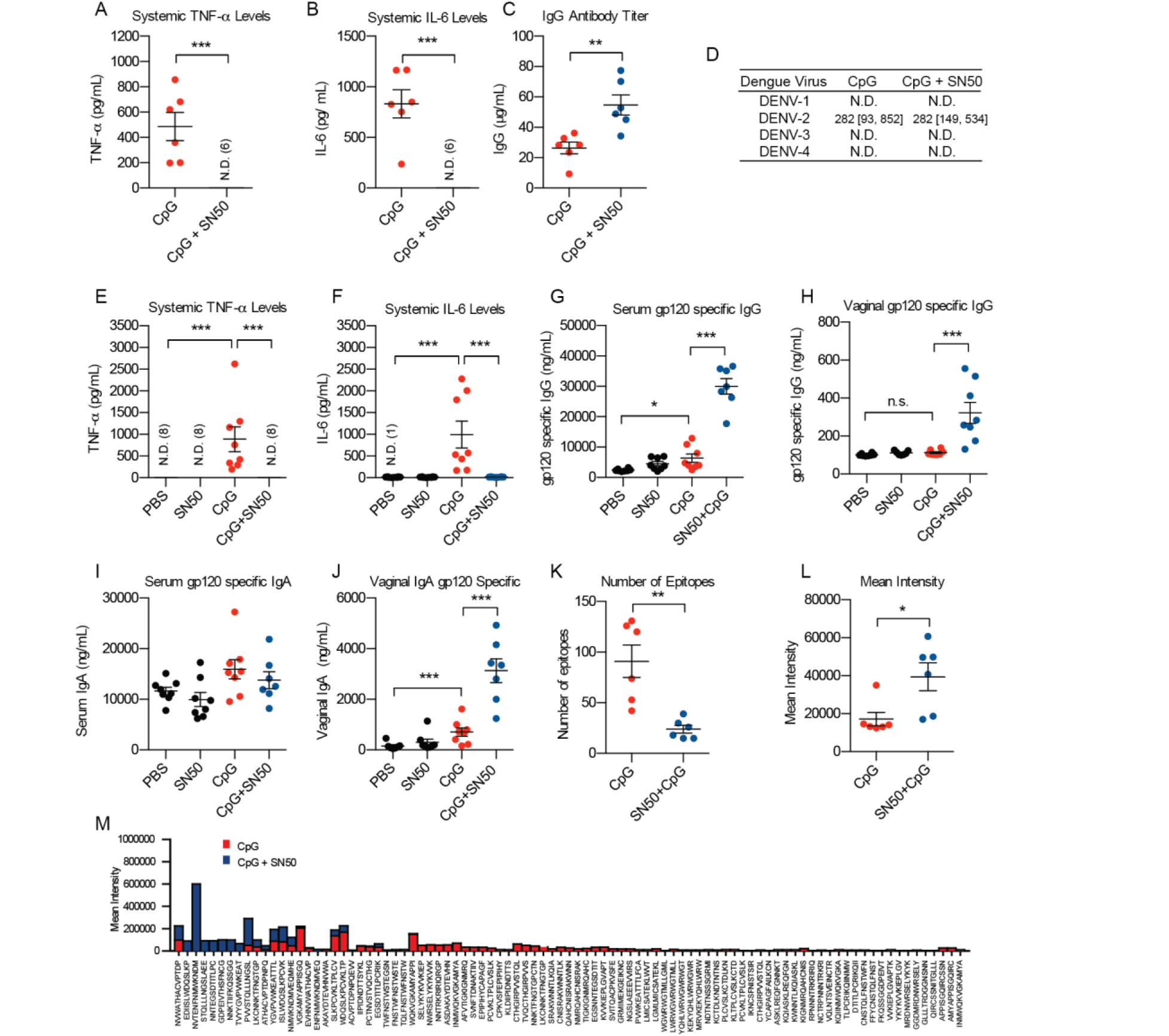
In vivo vaccination against dengue and HIV. (A) Systemic TNF-a levels 1h post-vaccination with DENV-2C antigen and CpG or CpG + SN50, n =6. (B) Systemic IL-6 levels 1h post-vaccination with DENV-2C antigen and CpG or CpG + SN50. (C) IgG antibody titer day 28 post vaccination with DENV-2C antigen. (D) Dengue virus neutralization. Geometric mean [95% confidence interval]. (E) Systemic TNF-a levels measured at 1h post-injection with gp120 and: PBS, CpG, SN50, SN50 + CpG, n = 8 (B) Systemic IL-6 levels measured at 1h post-injection with gp120 vaccinations (C) Serum anti-gp120 IgG antibody titer, day 28 after vaccination with gp120. (D) Vaginal anti-gp120 IgG antibody titer, day 28. (E) Serum anti-gp120 IgA antibody titer, day 28. (F) Vaginal anti-gp120 IgA antibody titer, day 28. (G) Number of g120 epitopes recognized by mice vaccinated with CpG or SN50 + CpG. (H) Mean intensity of recognized epitopes. (I) Mean intensity of each recognized epitope by CpG (red bars) or CpG + SN50 (blue bars).

To determine if SN50 alters the neutralization potential, we analyzed the neutralizing titer for four strains of dengue (**Fig. 3d**). We tested four serum samples against one strain representative of each dengue serotype. The differences in neutralization potential were not significantly different between the two groups implying that, similar to our flu results, SN50 improves the safety while maintaining the protective responses of vaccination.

## Analysis of influence of immune potentiator on HIV vaccination

To further test the efficacy of vaccines with SN50 and to attain a broader picture of the induced humoral immune response, we vaccinated mice with gp120, a viral coat protein from HIV necessary for infection and a target of many HIV vaccines, using CpG as the immune adjuvant. Mice vaccinated with CpG demonstrated high levels of both TNF-a and IL-6, whereas all other groups including mice vaccinated with CpG + SN50 demonstrated non-detectable levels of systemic cytokines at the 1h time point (**Fig. 3e, 3f**). On day 28, we analyzed antibody titers against gp120. The CpG + SN50 group induced a 4.7 fold higher anti-gp120 IgG antibody titer than the CpG group in the serum (**Fig. 3g, 3h**). This demonstrates that the addition of SN50 increases IgG antibody titer across multiple antigens and suggests that it may serve as a general immune potentiator. Because mucous membranes are particularly susceptible to HIV infection, we also measured the anti-gp120 IgG and IgA antibody titers in vaginal secretions (**Fig. 3i, 3j**). The CpG + SN50 group demonstrated a 4.4 fold increase in anti-gp120 IgA antibodies than mice vaccinated with CpG alone. These results suggest that SN50 with gp120 may help induce class-switching to IgA antibody isotype, while also enabling localization to the mucous membranes.

We next chose to determine if there were any alterations in the gp120 epitopes recognized by the resulting antibodies, using an overlapping peptide microarray. Interestingly, the number of epitopes recognized by CpG alone was higher than antibodies collected from CpG + SN50 mice; however, the fluorescent mean intensity of recognized epitopes is higher in the CpG + SN50 mice (**Fig. 3k, 3l**) – implying a higher concentration of antibodies against those epitopes. Upon closer inspection of particular epitopes recognized, we saw that adding the immune potentiator to the formulation shifts the epitope recognition, as different epitopes are recognized in the CpG alone and CpG + SN50, often exclusively in one condition or the other (**Fig. 3m**). This may prove valuable with diseases where the current recognized epitopes are not effective enough to provide protection. The most highly recognized epitope in the CpG + SN50 group corresponds to the epitope recognized by the recently isolated 35O22 monoclonal antibody (*46*). Antibodies isolated from mice vaccinated with CpG + SN50 also recognize the CD4 binding site recognized by several potent, broadly neutralizing antibodies (VRC 01, VRC03, b12). From these data we demonstrate that the addition of SN50 shifts the epitope selectivity in the case of gp120. Based on the epitopes recognized by the serum samples, we hypothesize that these antibodies may be more broadly neutralizing.

## Improvement of adjuvant responses across a variety of TLRs and Species

To examine the effects of the SN50 across a broad range of TLR agonists, we performed qPCR on RAW macrophages treated with SN50 followed by stimulation with agonists of different TLRs. We stimulated cells with SN50 and LPS (10 ng/mL), CpG (5 μg/mL), R848 (1 μg/mL) and Pam3CSK4 (100 ng/mL) and compared transcript levels to cells treated with TLR agonist alone (**Fig. 4a**). We chose these TLR agonists because they represent a subset of the compounds with promising potential for commercial use if the inflammatory side effects can be controlled. In RAW macrophages, we observed downregulation of TNF-a and IL-6 pro-inflammatory cytokine transcript levels. Across all agonists, the cell surface receptors CD86 and MHCII transcript levels were upregulated, compared to agonist alone, implying that cellular communication of the APC to the T cell may not be attenuated by the addition of SN50 and subsequent reduction in cytokine production.

**Figure 4.**
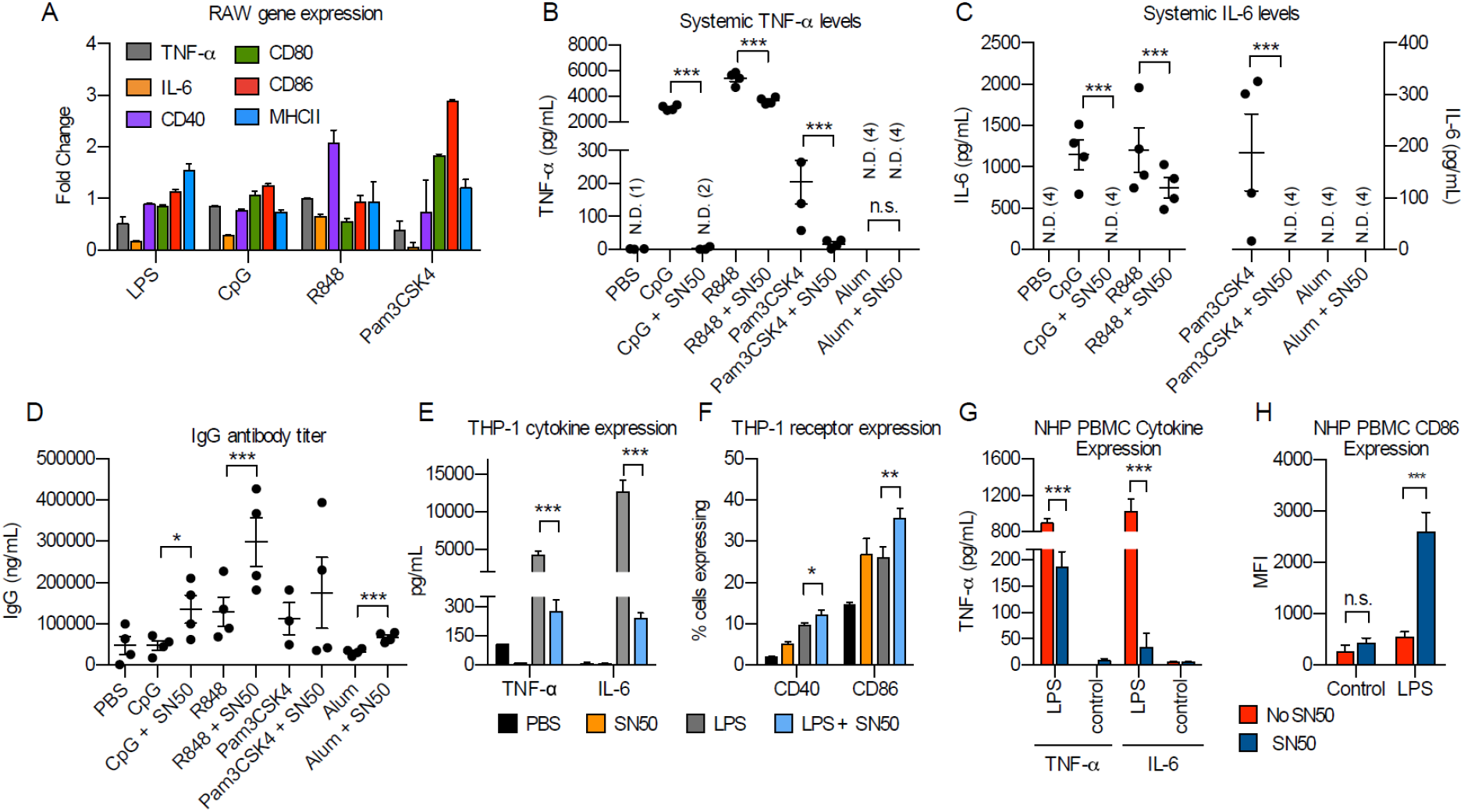
In vivo vaccinations across a broad range of adjuvants. (A) qPCR gene expression analysis of RAW macrophages stimulated with SN50 and TLR agonists compared to cells stimulated with TLR agonist alone. Pro-inflammatory cytokines TNF-a (grey bars) and IL-6 (orange bars) and cell surface receptors CD40 (purple bars), CD80 (green bars), CD86 (red bars) and MHCII (blue bars). (B) Systemic TNF-a cytokine levels of TNF-a measured at 1h post-injection with gp120 and: PBS, CpG, CpG + SN50, Pam3CSK4, Pam3CSK4 + SN50, R848, R848 + SN50, Alum, Alum + SN50, n =4. (C) Systemic IL-6 cytokine levels measured at 1h post-injection. (D) Serum IgG antibody titers, day 28. (E) Human THP-1 cell pro-inflammatory cytokines TNF-a and IL-6 in cell supernatant after treatment with PBS (black bars), SN50 (orange bars), LPS (grey bars), or LPS + SN50 (blue bars). (F) Cell surface receptor expression on human THP-1 cell after treatment with PBS (black bars), SN50 (orange bars), LPS (grey bars), or LPS + SN50 (blue bars). (G) Cytokine expression analysis of TNF-a and IL-6 in cell supernatant of NHP PBMCs 6h. No SN50 (red bars), SN50 (blue bars). LPS 1 μg /mL (H) CD86 expression of NHP PBMCs 18h. No SN50 (red bars), SN50 (blue bars).

To examine how this would translate in vivo, we vaccinated mice with CpG (50 μg), Pam3CSK4 (20 μg) and R848 (50 μg) using gp120 as the antigen. We chose to run these adjuvants alongside the most widely employed adjuvant, alum (250 μg).

With CpG, we observed complete elimination of systemic TNF-a and IL-6 proinflammatory cytokines (**Fig. 4b, 4c**). With R848 and Pam3CSK4 we saw a significant decrease in systemic cytokines. We hypothesize that SN50 is less effective at decreasing cytokines with R848 due to the low molecular weight of the molecule, enabling more rapid systemic distribution. Alum alone did not evoke a systemic cytokine response and the addition of SN50 did not alter the cytokine profile. The addition of SN50 increased the antibody titer for all adjuvants, including alum, demonstrating the broad potential use of this system to a large number of immune adjuvants (**Fig. 4d**).

To understand how this may translate to human vaccinations, we treated THP-1 monocytes with 1 μg /mL LPS with or without SN50. Cells treated with SN50 and LPS expressed dramatically lower levels of TNF-a and IL-6 (**Fig. 4e**). We also observed increased levels of CD40 and CD86 (**Fig. 4f**). Additionally, we examined the effects of SN50 on non-human primate rhesus macaque (NHP) primary peripheral blood mononuclear cells (PBMCs). We stimulated NHP PBMCs with SN50 and LPS or LPS alone for 6h and analyzed the cell supernatant for pro-inflammatory cytokines. Cells stimulated with LPS demonstrated high levels of TNF-a and IL-6 in the cell supernatant, cells with SN50 demonstrated significant reduction in cytokine levels (**Fig. 4g**). We observed that CD86 expression was upregulated 2-fold in cells stimulated with SN50 and LPS compared to cells stimulated with LPS alone (**Fig. 4h**). This implies that SN50 may work similarly in NHP and humans as it does in mice.

## CONCLUSION

Using a broad range of TLR agonists, we show both in vitro and in vivo that a cell permeable inhibitor of the p50 subunit of NF-kB, potentiates the immune response – reducing inflammation while increasing antibody responses. Co-administration of CpG with the immune potentiator results in significantly reduced levels of proinflammatory cytokines, often at undetectable levels. At the same time, this reduction in inflammation results in a 3-fold increase in the IgG titer of antibodies for the model antigen OVA. We examined how potentiation would enhance the capabilities of the adjuvants to improve the immune response. In our influenza model we directly examined side effects in response to the current commercial flu vaccine and Fz + CpG and determine that adding SN50 reduces side effects and systemic pro-inflammatory cytokine levels. We also demonstrate that the safety profile can be enhanced without negatively effecting the protective response. After vaccinated mice were challenged with influenza A, mice with SN50 added to the vaccine, lead to increased survival, less weight loss and less change in body temperature. To study the effects of potentiation on TLR agonists as vaccine adjuvants, we selected three diseases – influenza, dengue and HIV – all of which have had different challenges in vaccine development. In dengue vaccination, the goal is to increase antibody neutralization potential while maintaining a safe profile. We demonstrate that there are no detrimental effects to dengue neutralization of antibodies with SN50, enabling us to mitigate side effects but maintain the protective response. In HIV, we vaccinated with HIV envelope protein gp120, CpG and SN50, increased both IgG and IgA titers. This method appears quite general as it works with many TLR agonists and antigens. SN50 is one of hundreds of similar NF-kB inhibitors. When used in combination with the appropriate TLR agonist, many may prove useful for eliciting specific and potentially tunable responses for distinct vaccines or immunotherapies. This methodology may find use in reducing the systemic side effects associated with inflammation seen in many adjuvanted vaccines (*19*). This method has the potential to enable a variety of PRR agonists to be used safely in vaccines, increasing the diversity of adaptive immune profiles and widening the scope of disease prevention and treatment.

In conclusion, we have demonstrated that using a specific NF-kB inhibitor in combination with common immune adjuvants can decrease pro-inflammatory cytokine production while boosting cell-surface receptor expression for effective antigen presentation and T cell activation in mouse, human and NHP primary cells. The use of this inhibitor in vivo completely reduced systemic TNF-a and IL-6 to baseline levels while increasing the downstream adaptive humoral response from the vaccination. These phenomena were observed across a broad range of antigens for a variety of pathogens demonstrating that this may prove a general strategy for improving vaccination response while conforming to strict safety standards. There are hundreds of documented immune adjuvants that provide adequate protection against diseases but induce unsafe levels of inflammation to be approved for clinical use (7,45). Additionally there are hundreds of NF-kB inhibitors, some already with FDA approval, that could be multiplexed with different TLR agonists to provide a broad range of responses (44). We anticipate this framework will enable a variety of TLR agonists to be used safely in human vaccines, increasing the diversity of adaptive immune profiles and widening the scope of disease prevention and treatment.

## Supporting information

## Acknowledgments

The authors thank Prof. Amanda Burkhardt and Adrienne Gilkes for training and assistance with in vivo techniques. The authors thank Rie Nakajima for assistance imaging microarrays and Dan Piraner for help with image analysis. The authors thank Alfred Chon for assistance with HPLC optimization and Naorem Nihesh for training and assistance with peptide synthesis. The authors acknowledge Dr. Jin Qin at the University of Chicago for training and assistance with mass spectrometry.

## Funding

We would like to acknowledge support by the NIH (1U01Al124286-01 and 1DP2Al112194-01, GM099594). Prof. Esser-Kahn thanks the Pew Scholars Program and the Cottrell Scholars Program for generous support. B.A.M. thanks NSF-GRFP (DGE-1321846). We would like to thank NSF instrumentation grant CHE-1048528. This work was supported, in part, by a grant from the Alfred P. Sloan foundation.

## Author contributions

B.A.M. and A.E.K conceived of and designed the project and experiments and wrote the manuscript. B.A.M., R.S., Y.E.B, D.B., K.B., B.G.M. performed experiments. B.A.M. and S.Y. synthesized materials.

## Competing interests

The authors report no competing interests.

## Data and materials availability

All data are available in the main text or the supplementary materials.

## Supplementary Materials

Materials and Methods Figures S1-S9

