## Supplementary material for "Immune potentiator for increased safety and improved protection of vaccines by NF-kB modulation"

**This PDF file includes:**

Materials and Methods
Figs. S1 to S9

### Materials and Methods

#### RAW Blue NF-kB Assay

RAW-Blue™ NF-kB cells (Invivogen) were passaged and plated in a 96 well plate at 100k cells/ well in 180 µL DMEM containing 10% HIFBS. Cells were incubated at 37 °C and 5% $CO_2$ for 1 h. SN50 was added at indicated concentrations, cells were incubated 1 h. Immune agonists were added at their indicated concentrations. The volume of each well was brought to 200 µL and incubated at 37 °C and 5% $CO_2$ for 18 h. After 18 h, 20 µL of the cell supernatant was placed in 180 µL freshly prepared QuantiBlue (Invivogen) solution and incubated at 37 °C and 5% $CO_2$ for up to 2 h. The plate was analyzed every hour using a Multiskan FC plate reader (Thermo Scientific) and absorbance was measured at 620 nm.

#### THP-1 Blue NF-kB Assay

THP-Blue™ NF-kB cells (Invivogen) were passaged and plated in a 96 well plate at 400k cells/ well in 180 µL RPMI 1680 containing 10% HIFBS. Cells were incubated at 37 °C and 5% $CO_2$ for 1 h. SN50 was added at indicated concentrations and cells were incubated for 1 h. Immune agonists were added at their indicated concentrations. The volume of each well was brought to 200 µL and incubated at 37 °C and 5% $CO_2$ for 18 h. After 18 h, the plate was spun down at 400 x g (Allegra X-30, Beckman Coulter) and 20 µL of the cell supernatant was placed in 180 µL freshly prepared QuantiBlue (Invivogen) solution and incubated at 37 °C and 5% $CO_2$ for up to 2 h. The plate was analyzed every hour using a Multiskan FC plate reader (Thermo Scientific) and absorbance was measured at 620 nm.

#### Gene expression

RAW 264.7 macrophages or THP-1 cells were passaged and plated in a cell culture treated 6- well plate at 4 $x10^6$ cells/ well in 1.5 mL DMEM or RPMI (respectively) containing 10% HIFBS. SN50 (250 µg/mL) or PBS was added to wells and cells were incubated for 1 h at 37 °C and 5% $CO_2$ for 6 h. RNA was extracted using RNeasy Plus Mini kit (Qiagen). RT-PCR was performed using RT2 first strand kit (Qiagen) and BioRad thermocycler according to manufacturer's protocol. cDNA was stored at -20 °C. RT2 SYBR ROX qPCR Master mix (Qiagen) was used according to manufacturer's protocol. qPCR amplification was performed using a Stratagene Mx3005P thermocycler.

#### Intracellular cytokine staining

#### BMDCs

Monocytes were harvested from 6-week-old C57BL/6 mice. Monocytes were differentiated into dendritic cells (BMDCs) using supplemented culture medium: RPMI 1640 (Life Technologies), 10% HIFBS (Sigma), 20 ng/mL granulocyte-macrophage colony stimulating factor (produced using "66" cell line), 2 mM Lglutamine (Life Technologies), 1% antibiotic-antimycotic (Life Technologies), and 50 µM beta-mercaptoethanol (Sigma). After 5 days of culture, BMDCs were incubated with 250 µg/mL SN50. After 1 h CpG ODN 1826 (IDT) and 1 µL/mL GolgiPlug (BD Biosciences) was added. Cells were incubated for 6 h at 37 °C in and 5% CO2. Cells were stained for CD40 (Biologend, 124610), CD86 (Biolegend, 105109), and intracellular IL-6 (Biolegend, 504508) and TNF-a (Biolegend, 506308) cytokine production and analyzed using BD Accuri C6 flow cytometer.

#### SEM analysis

Sample suspensions obtained directly from injection mixtures were dried for 24 h, mounted on carbon tape, and sputter coated (South Bay Technologies) with approximately 2-4 nm of Au/Pd

60:40 or Ir. Scanning electron microscopy (SEM) of the sample suspensions was performed using an FEI Quanta 3D FEG dual beam (SEM/FIB) equipped with Inca EDS (Oxford Instruments).

**In vivo studies**

**Animals**

All animal procedures were performed under a protocol approved by the University of Chicago Institutional Animal Care and Use Committee (IACUC). 6-8 week-old C57/B6 female mice were purchased from Jackson Laboratory (JAX). All compounds were tested for endotoxin prior to use. All vaccinations were administered intramuscularly in the hind leg. Blood was collected from the sapheneous vein at time points indicated.

Antigens were purchased from Sino Biological (HIV subgroup M, Influenza A H1N1 (A/California/04/2009) Hemagglutinin / HA Protein, Dengue virus DENV-2 (Strain New Guinea C) Capsid protein / DENV-C Protein (His Tag), Virogen (HIV-1 env (gp41) antigen) or Invitrogen (Vaccigrade Ovalbumin). Vaccigrade CpG ODN 1826 was purchased from Invivogen or Adipogen. SN50 was synthesized via solid phase peptide synthesis as previously described and purified using Gilson preparatory HPLC.

**Vaccinations**

Mice were anesthetized lightly with isoflurane and injected intramuscularly in the hind leg with 50 uL containing antigen, adjuvant and PBS. Antigen doses: ovalbumin (100 µg), DENV2-C (5 µg) and gp120 (3 ug). CpG dose, 50 µg. SN50, 500 µg. TNF-aN, 30 µg. IL-6N, 30 µg.

**Plasma cytokine analysis**

Blood was collected from mice at specified time points in 0.2 mL heparin coated collection tubes (VWR Scientific). Serum was isolated via centrifugation 2000 x g for 5 min. Supernatant was collected and stored at -80 °C until use. Serum was analyzed using BD Cytometric Bead Array Mouse Th1/Th2/Th17 cytokine kit or Mouse Inflammation cytokine kit according to manufacter's protocol. Briefly, beads containing antibodies for desired cytokines were mixed with 50 µL serum and 50 µL PE detection reagent and incubated for 2 h. Beads were washed and analyzed using BD Accuri C6 flow cytometer. Data was analyzed using BD Accuri C6 software and Graphpad Prism.

**Antibody titer analysis**

Mice were vaccinated with indicated formulations. Blood was collected at time points indicated in 0.2 mL heparin coated collection tubes (VWR Scientific) for plasma or uncoated tubes for serum. Plasma was isolated via centrifugation (2000 x g, 5 min). Serum was isolated by allowing blood to clot for 15- 30 min RT and centrifuging (2000 x g for 10 min) at 4 °C. Serum was analyzed using a quantitative anti-ovalbumin total Ig's ELISA kit (Alpha Diagnostic International) according to the specified protocol. Total IgG and IgA was analyzed using total mouse IgG or IgA uncoated ELISA (Invitrogen) and was analyzed using Multiskan FC plate reader (Thermo Scientific) and absorbance was measured at 450 nm. Data was analyzed using Graphpad Prism.

**Influenza Challenge Model**

**Animals**

All animal procedures were performed under a protocol approved by the Illinois Institute of Technology Research Institute (IITRI) Animal Care and Use Committee (IACUC). 6-8 week-old C57/B6 female mice were purchased from Charles River. All compounds were tested for endotoxin prior to use. All vaccinations were administered intramuscularly in the hind leg.

Initial group assignments were assigned to using a computerized randomization procedure based on body weights that produce similar group mean values [ToxData® version 3.0 (PDS Pathology Data Systems, Inc., Basel, Switzerland)]. Mice were vaccinated by i.m. injection into the hind leg on Days 0 and 21. The vaccine material used in this study is Fluzone® quadrivalent influenza vaccine (Sanofi Pasteur). Each 0.5 mL dose of Fluzone® contains at least 15 μg of hemagglutinin (HA) from each of the following four influenza strains recommended for the 2017/2018 influenza season: A/Michigan/45/2015 X-275 (H1N1)pdm09-like strain, A/Hong Kong/4801/2014 X-263B (H3N2)-like strain, B/Phuket/3073/2013-like strain and B/Brisbane/60/2008-like strain. At least 1 μg of each strain was used in vaccination of the mice. Body weights were collected 24 hr, 48 hr and 72 hr post-prime vaccination. Body temperatures were collected 1 hr, 3 hr, 24 hr, 48 hr and 72 hr post-prime vaccination. Blood samples were collected on days 0, 14, 28, 42, 56. Plasma was collected on day 0. Serum was collected on days 14, 28, 52 and 56. Five animals from each group were humanely euthanized on day 14 post-vaccination. Spleens were collected for T cell analysis. On day 43 post-vaccination, all mice were challenged via intranasal route with a lethal dose of A/Michigan/45/2015. The dose level of challenge virus used was an equivalent of 5 LD50. For inoculation, mice were anesthetized with a ketamine (80 mg/kg) and xylazine (10 mg/kg) mixture. Once anesthetized, 0.025 mL of inoculum was delivered dropwise into the nares. The mouse was held upright to allow the virus to be inhaled thoroughly then returned to its cage. After challenge, body weights and temperature readings were recorded daily through a transponder (BioMedic data systems, Seaford, DE) implanted subcutaneously in each mouse. Animals were monitored for morbidity/mortality for 14 days post-infection. Any animals meeting pre-determined moribund criteria (>20% weight loss) were humanely euthanized. Three animals from each group were humanely euthanized on day 3 post-challenge (Day 45) and lungs collected for viral quantitation by plaque assay/TCID50. Tissues for viral titers were weighed then flash frozen in an ethanol/dry ice bath or liquid nitrogen and stored at ≤-65°C. Frozen organs were thawed at 37 °C for 5 min. Once thawed, organs were homogenized in MEM 10% w/v using a Bead Ruptor 12 (Omni International, Kennesaw, Georgia) in tubes containing 1.4 mm ceramic beads. Homogenized organs were centrifuged at 2,000 x g for 5 min to remove cellular debris. The resulting supernatant was serially diluted 10-fold then transferred into respective wells of a 96-well plate containing a monolayer of Madin-Darby Canine Kidney Cells (MDCK) cells for titration. The TCID50 assay will be performed. TCID50 titers will be calculated using the method of Reed-Meunch. The remaining 5 mice in each group were monitored for the remaining days of the challenge.

**Neutralization assays**

Serum samples were tested against a representative of each dengue serotype (DENV-1: strain Hawaii; DENV-2 strain New Guinea C; DENV-3 strain Philippines/H87/1956 and DENV-4 strain H241). Sera was serially diluted two-fold, (starting dilution 1:100) then incubated with standardized virus concentration of 50-120 PFU of each strain. The serum:virus mixture was transferred into respective wells of a 96-well plate which contained a monolayer of Vero cells. The cells were incubated for 40 hours at 37 °C. After 40 hours of incubation, the cells were fixed with 1.0% paraformaldehyde and stained by Anti-Flavivirus Group Antigen Antibody, clone D1-4G2-4-15 (Millipore Billerica, MA) followed by peroxidase-conjugated goat anti-mouse IgG (Kirkegaard and Perry Laboratories, Gaithersburg, MD). Spots were developed using TrueBlue Peroxidase Substrate (Kirkegaard and Perry Laboratories, Gaithersburg, MD). Plaques were visualized and counted using an ELISPOT instrument. Plaque reduction neutralization test titers (PRNT) were expressed in terms of conventional 50% PRNT end-point titers.

**T cell analysis**

Spleens were harvested from mice as described above at time point indicated. Splenocytes were isolated by pressing spleen fragments through a strainer attached to a 50-mL conical tube using a syringe plunger. Cells were washed through the strainer with PBS and centrifuged at 500 x g for 10 min. Supernatant was aspirated and the pellet was resuspended in 2 mL of pre-warmed lysing solution (BD Pharm Lyse™ lysing solution) and incubated at 37 °C for 2 minutes. 30 mL of PBS was added and cell suspension was centrifuged at 500 x g for 10 minutes. Supernatant was discarded and cells were resuspended in RPMI containing 10% HIFBS at 2 x $10^6$ cells/ mL. 500 µL was added to 24 well plate. Cells were incubated with 10 µg/ mL Influenza A H1N1 (A/Michigan/45/2015) Hemagglutinin / HA1 Protein (His Tag) (SinoBiological). After 1h GolgiStop (BD Biosciences) was added and cells were incubated for 11 h at 37 °C and 5% $CO_2$. Cells were centrifuged 500 x g for 10 min and stained for CD4/ IL-4 (Biolegend o FITC anti-mouse CD4 [RM4-5], PerCP/Cy5.5 anti-mouse IL-4 [11B11]) or CD8/ IFN-y (FITC anti-mouse CD8a [53-6.7], PE anti-mouse IFN-γ [XMG1.2]) using BD Cytofix/Cytoperm Fixation/Permeabilization Solution Kit according to manufacter's protocol and analyzed using a NovoCyte flow cytometer (ACEA Biosciences, Inc.).

**Epitope analysis**

Mouse serum was collected as described above and samples were analyzed using Multiwell RepliTope™ microarray for appropriate antigen (JPT Innovative Peptide Solutions) according to manufacterer's protocol. Briefly, serum samples were diluted in 3% BSA in 1x TBS-Buffer + 0.1% Tween20 (TBS-T) to a final concentration of 10 µg/mL. The microarray was fitted with an ArraySlide 24-4 chamber (JPT Innovative Peptide Solutions) to enable multi-sample analysis. 150 uL diluted serum was added to samples wells and incubated for 1h at 30 °C. Wells were washed 5x with TBS-T. 150 µL secondary antibody (1 µg/mL) was added to wells and incubated RT for 1h. Wells were washed 5x with TBS-T and 2x with nanopure water. Arrays were imaged using a Caliber I.D. RS-G4 confocal microscope and analyzed using ImageJ.

**Safety and protection score**

We assigned a safety score comprised of systemic TNF-a, IL-6 levels and weight loss post-vaccination. A score for each TNF-a, IL-6 and weight loss was assigned for each mouse. The safety score of a single mouse represents the summation of these individual scores. A protection score was assigned based on survival, change in body weight and change in body temperature post-challenge. Scores were determined by dividing values into quartiles, and assigned a number 0 to 4 based on the quartile. Higher values indicate an improved safety profile (lower TNF-a or IL-6, less weight loss after vaccination) or improved protection (survival, less weight loss, higher body temperature after challenge).

**Statistics and replicates**

Data is plotted and reported in the text as the mean ± s.e.m. Sample size is as indicated in biological replicates in all in vivo and in vitro experiments. The sample sizes were chosen based on preliminary experiments or literature precedent indicating that the number would be sufficient to detect significant differences in mean values should they exist. P values were calculated using a two-tailed unpaired heteroscedastic t-test.

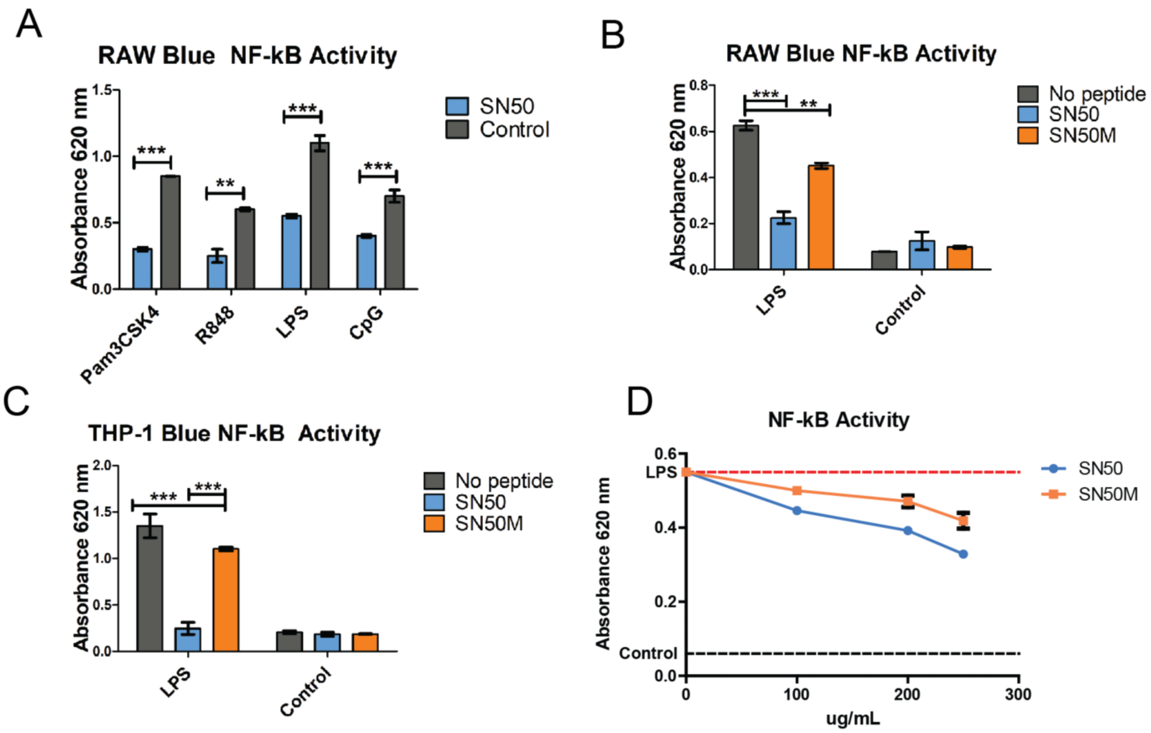


**Figure S1. NF-kB activity in mouse and human cells**. (A) NF-kB activity in RAW blue macrophages stimulated with various TLR agonists and SN50 (blue bars), TLR agonists alone (grey bars). (B) RAW blue NF-kB activity of cells stimulated with SN50 (blue bars) and the control peptide SN50M (grey bars). (C) THP-1 NF-kB activity of cells stimulated with no peptide (grey bars), SN50 (blue bars) or SN50M (orange bars). (D) Concentration screen of 100 ng/ mL LPS and various concentrations of SN50 (blue line) and SN50M (orange line).

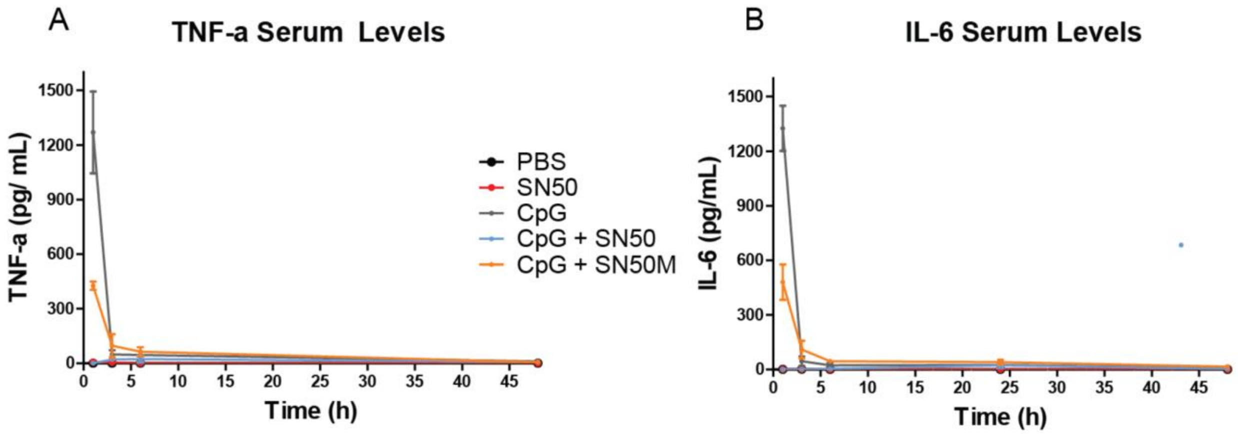


**Figure S2. Proinflammatory Cytokine Analysis Time Course** (A) Systemic TNF-a levels measured at 1h, 3h, 6h, 24, 48 h post vaccination. (B) Systemic IL-6 levels.

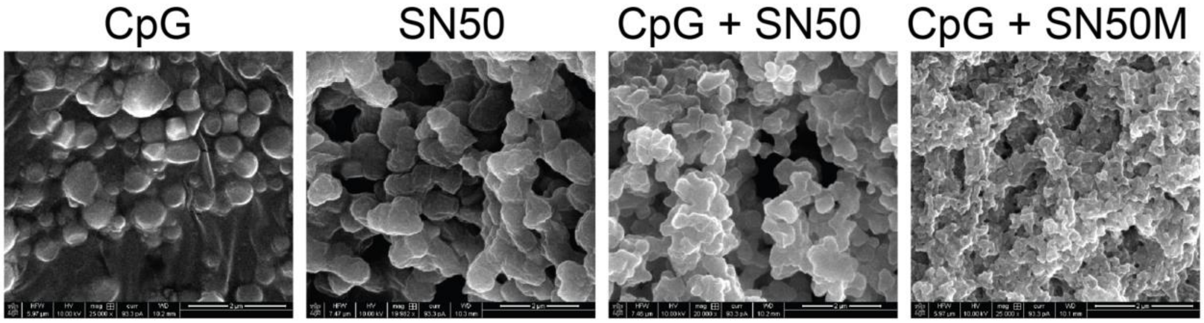


**Figure S3. SEM image of OVA vaccinations.** (A) CpG + OVA (B) SN50 + OVA (C) CpG + SN50 + OVA (D) CpG + SN50M + OVA. Scale bar 2 um.

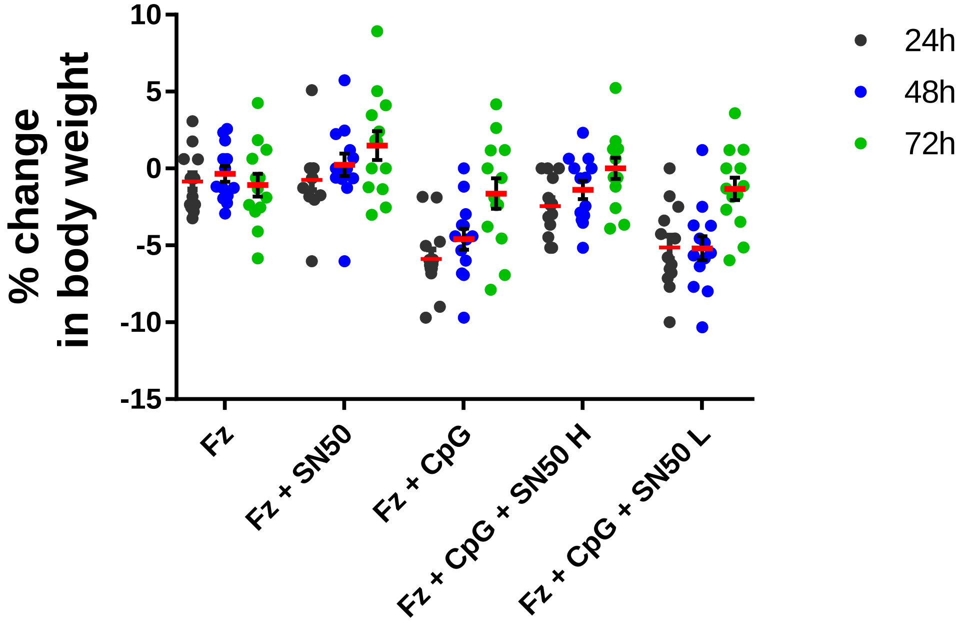


**Figure S4. Weight Loss Post-Vaccination.** Weight loss 24h (black dot), 48h (blue dot) and 72h (green dot) post prime. Fz = Fluzone 2017-2018 flu vaccine.

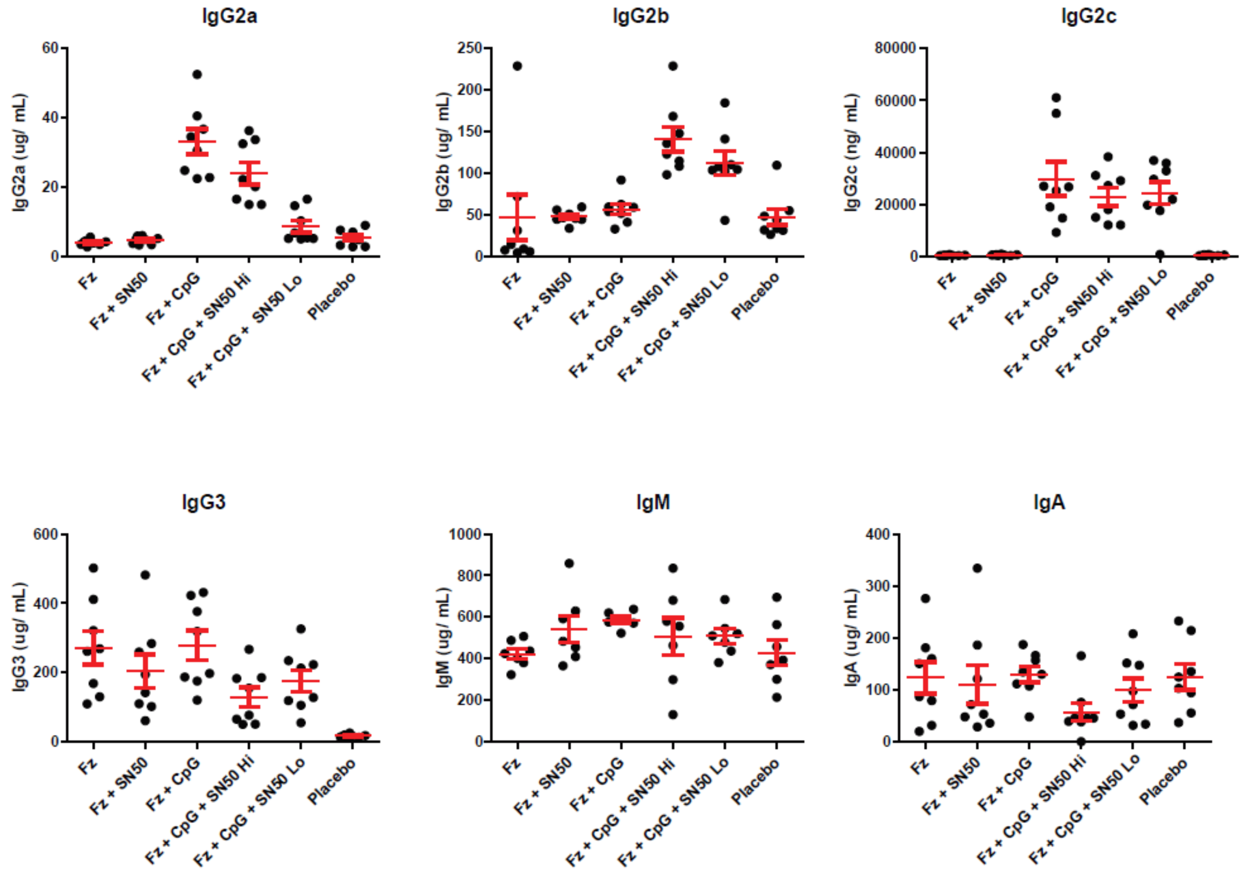


**Figure S5. Day 28 Antibody Titers.** Fz = Fluzone 2017-2018 flu vaccine.

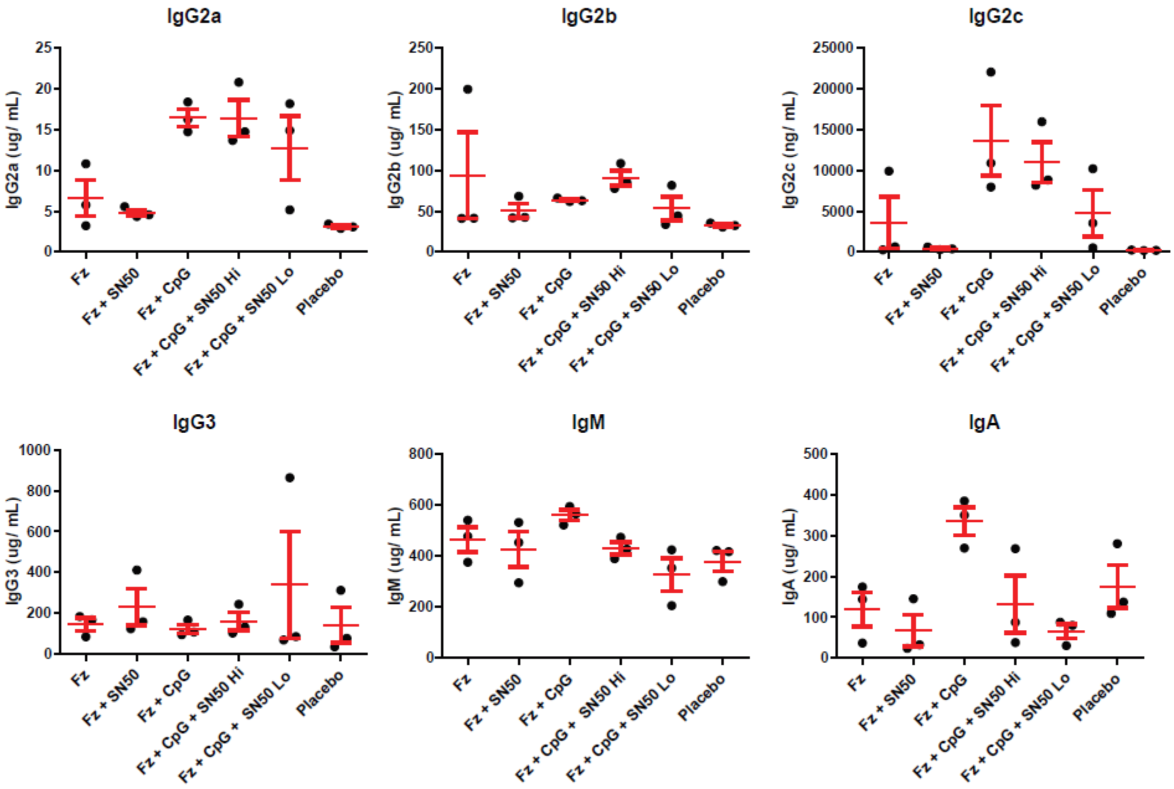


**Figure S6. Day 46 Antibody Titers (d3 post infection).** Fz = Fluzone 2017-2018 flu vaccine.

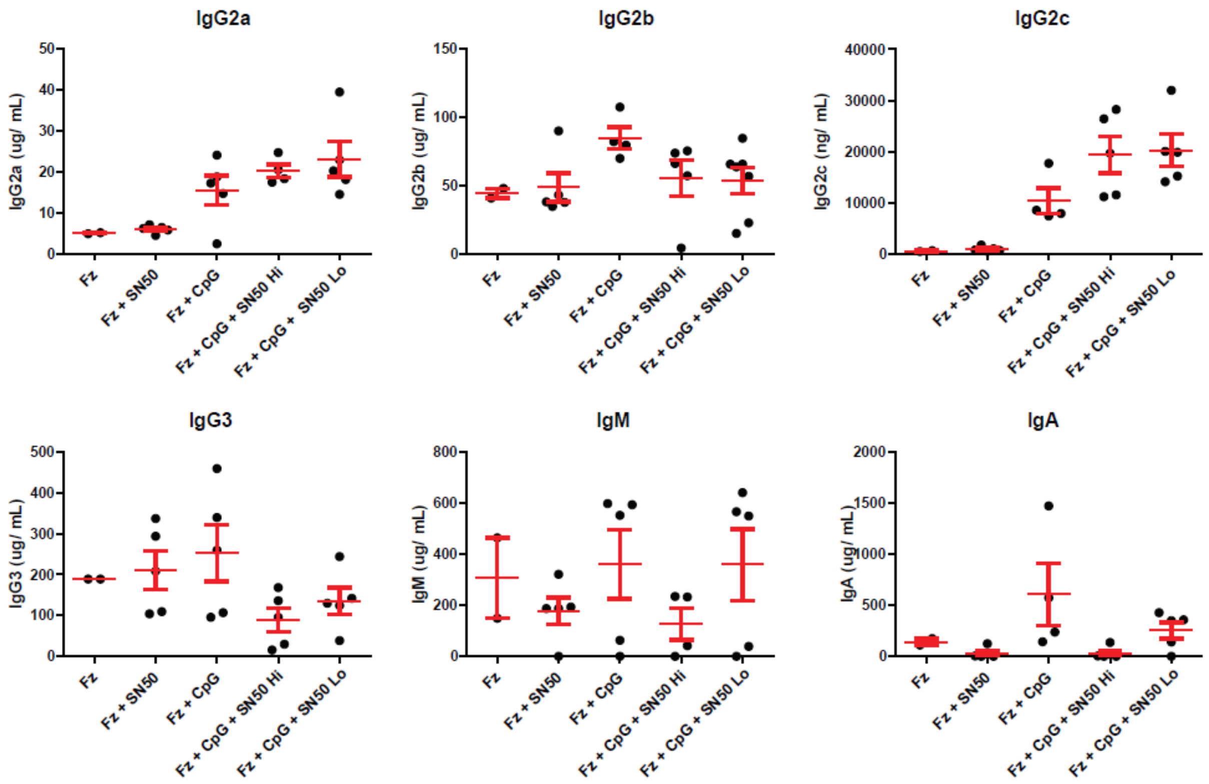


**Figure S7. Day 57 Antibody Titers of surviving mice (d14 post infection).** Fz (2), Fz + SN50 (5), Fz + CpG (5), Fz + CpG + SN50 Hi (5), Fz + CpG + SN50 Lo (5). Fz = Fluzone 2017-2018 flu vaccine.

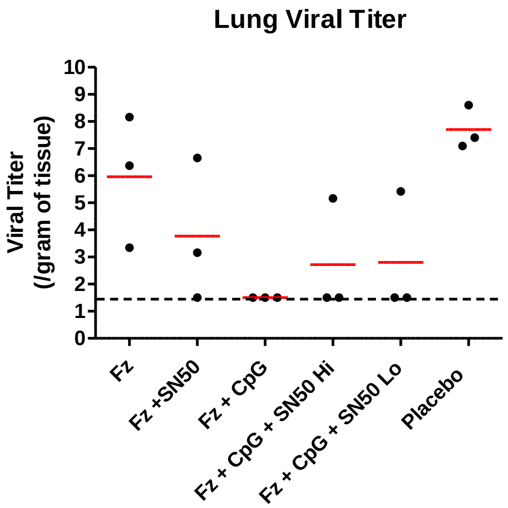


**Fig. S8. Lung viral titer d3 post-infection.** Fz = Fluzone 2017-2018 flu vaccine.

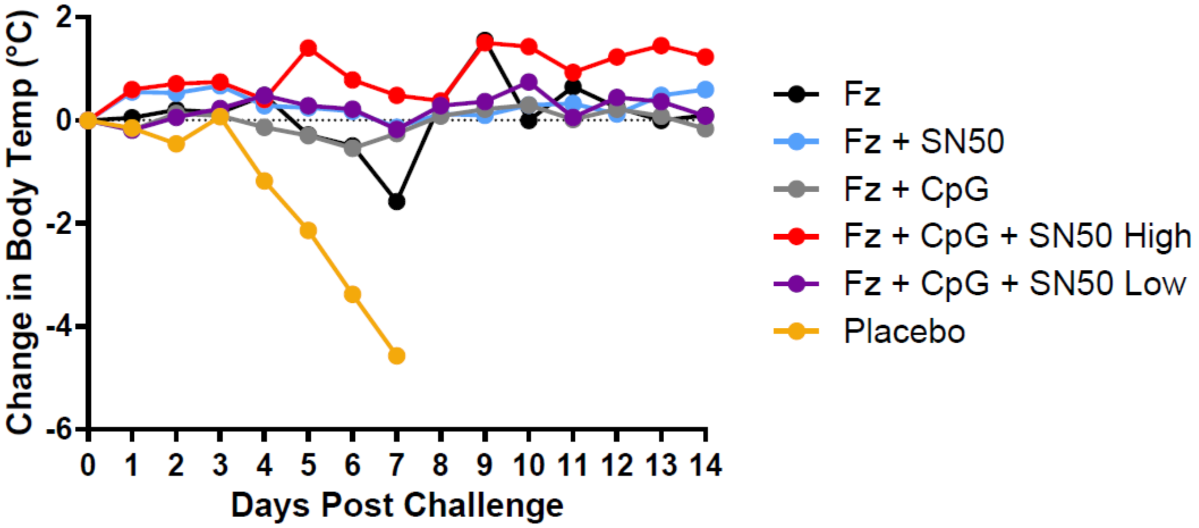


**Fig. S9. Full temperature curve for 14 days post-challenge.** Fz (black line), Fz + SN50 (blue line), Fz + CpG (grey line), Fz + CpG + SN50 Hi (red line), Fz + CpG + SN50 Lo (purple line), Placebo (yellow line).
